# Potato cultivars use different root physiological and molecular mechanisms to acclimate to salt stress

**DOI:** 10.1101/2024.07.05.602205

**Authors:** Michael Nicolas, Jort Bouma, Jan Henk Venema, Hanneke van der Schoot, Francel Verstappen, Thijs de Zeeuw, Sanne E. Langedijk, Damian Boer, Johan Bucher, Marten Staal, Ben Krom, J. Theo M. Elzenga, Richard G.F. Visser, Christa Testerink, Rumyana Karlova

## Abstract

- Soil salinity induces osmotic stress and ion toxicity in plants, detrimentally affecting their growth and development. Potato (*Solanum tuberosum*) faces yield reductions due to salt stress. The mechanisms of salt stress resilience, especially in adventitious roots, remain unknown.
- We investigated the resilience of three potato cultivars - Desirée, Innovator, and Mozart - by studying their physiological and transcriptomic responses to salt stress.
- Our findings reveal that under salt stress, the growth of stolons and stolon node roots is similarly reduced unlike tubers, even though they are physically connected. Surprisingly, tubers accumulate Cl^-^ but not Na^+^ under salt stress, suggesting an active Na^+^ exclusion mechanism. Innovator showed the lowest suberin and lignin deposition before salt stress and higher K^+^ leakage, leading to a stronger initial stress response with high ABA content and a distinct transcriptomic pattern. Nevertheless, Innovator was the most resilient, displaying lower growth, salt-tolerance index and tuber yield reduction. Transcriptomic analysis revealed several K^+^/Na^+^ channel genes which might regulate ions homeostasis during salt stress, in particular in Innovator.
- Altogether, we conclude that acclimation ability, rather than initial protection of roots against salt, prevails in long term salt-stress resilience of potato.

## Introduction

Soil salinity is worldwide problem that reduces crop biomass and yield, causes early senescence, and can lead to plant death (Ghosh et al., 2001; Jaarsma et al., 2013; Van Zelm et al., 2020). Factors contributing to salinisation include soil erosion, poor irrigation, drought, flooding and sea level rise. About 10% of arable soil and 25 to 30% of irrigated land are saline worldwide, and situation is expected to worsen with climate change which threats food security as the population grows (Dasgupta et al., 2009; Nawaz et al., 2010; Shahid et al., 2018; Dahal et al., 2019; Ivushkin et al., 2019; Hassani et al., 2020).

Salt impacts plant growth through osmotic and ion toxicity stresses. Different phases can be distinguished from the early Na^+^ sensing to growth recovery. Early Na^+^ accumulation in root cells stimulates different short- and long-range Ca^2+^ waves, which, activate the SALT OVERLY SENSITIVE (SOS) pathway (Lamers et al., 2020). In this early response HKT1 channel is also activated to exclude Na^+^ from the cell when NHXs ion transporter sequesters it in the vacuole (Van Zelm et al., 2020). The remaining cytosolic Na^+^ physically competes with K^+^ but does not achieve its function, hence impacting intracellular K^+^ homeostasis. Plant cell regulates the Na^+^/K^+^ ratio according to Na^+^ amount to maintain cellular osmotic and turgor pressure (Duan et al., 2013; Zou et al., 2022). This early sensing phase is followed in the next few hours by a stop and quiescent phases where auxin, ABA and ethylene signalling play a major role, allowing cell transcriptional reprogramming for acclimation. This transcriptomic salt stress response allow the initiation of the growth recovery phase where gibberellins, brassinosteroids, and jasmonic acid usually play a crucial role (Van Zelm et al., 2020). Salt stress response also includes an increased regulation of apoplastic transport to the stele by a higher lignin deposition around endodermal cells, the Casparian strip, which involves several MYB transcription factors (Calvo-Polanco et al., 2021; Li et al., 2023). Likewise, a stronger suberin deposition in the cell wall decreases Na^+^ permeability (Ranathunge and Schreiber, 2011; Barberon et al., 2016). This new growth initiation requires photosynthetic carbon influx from the leaf, the source tissue (Ho, 1988; Sonnewald and Fernie, 2018). By reducing growth in sink organs, salt stress decreases sugar unloading and hence sink strength which can lead to carbon starvation in case of severe salt stress (Smith and Stitt, 2007).

Potato (*Solanum tuberosum*) is the fourth major crop (FAO, 2017) and represents an important species for food security worldwide. Potato is a moderately salt-sensitive crop, exhibiting reduced tuber yield under saline conditions (Nawaz et al., 2010; Jaarsma and de Boer, 2018). Previous research identified cv. Innovator as a more tolerant cultivar in an *in vitro* assay, showing a lower growth reduction of shoot, roots and tubers at different salt concentrations (Ahmed et al., 2020), while explants of cv. Desirée were found to be more tolerant than cv. Mozart, displaying a higher vegetative growth and accumulating more proline and less H_2_O_2_ in hydroponics (Jaarsma et al., 2013). In addition to the traditional development of shoots, branches, and leaves, potato plant develops different peculiar belowground organs, namely stolons, tubers and adventitious roots. Stolons are underground branches from which the tubers emerge. Tuber formation occurs in the meristematic regions where sucrose is converted into glucose 6-phosphate, which is subsequently transported into the amyloplasts to be transformed into starch (Ewing and Struik, 1992). Potato plants usually develop from tubers, not seeds, and only produce adventitious roots. Three different kinds of adventitious roots can be distinguished (Kratzke and Palta, 1985); the basal root, emerging from the base of the shoot underground; the stolon root, which grow from the belowground shoot node along with the stolon; and the stolon-node root, developing from the stolon nodes. Very little is known about how salt stress affects belowground organs of potato and in particular its adventitious roots.

Here, based on the literature, we selected Innovator, Desirée, and Mozart as contrasting cultivars to further study their mechanisms of salt stress tolerance. Our data showed that Innovator has a lower yield reduction in salt stress conditions, which however, is linked with a higher water loss compared to Desirée. Interestingly, we found that tubers actively exclude Na^+^ ions but do accumulate K^+^ and Cl^-^ ions. Innovator is initially less protected against salt stress by displaying the lowest suberin and lignin depositions prior salt stress. However, it shows the strongest acclimation potential by adequately accumulating suberin and lignin and by reducing K^+^ leakage in roots, after experiencing a significant stress associated with a massive ABA accumulation in the roots. This suggests that the fast suberin deposition under salt stress but not the initial amount correlates with the degree of resilience of the cultivars. RNA-seq transcriptomic analysis showed that the dynamics of salt stress response in roots are also different in Innovator. We identified several transcription factor families that temporally regulate the salt stress response in potato roots.

## Materials and Methods (word counts: 1084)

### Plant materials and growth conditions

*S. tuberosum* L. cvs. Desirée, Innovator and Mozart were propagated *in vitro* from single-node stem cuttings on Murashige and Skoog medium including vitamins (Duchefa Biochemie BV, Haarlem, the Netherlands) containing 2% sucrose and 0.6% plant agar, 0.05% MES (Duchefa Biochemie BV) in chambers at 24 °C in an LD photoperiod (16 h light, 8 h dark). Five-node plants were cut at the base and transferred to a new box during seven days for rooting (greenhouse experiment at Unifarm WUR conditions) or 3 days for the hydroponic experiment.

### Plant phenotyping

Plants were transferred in 2l pot containing vermiculite (Agra-vermiculite no.3, Helza-Hobbyzaden). Growing conditions were 16/8 h light/dark with 21°C/18°C day/night temperature and 60% humidity. Plants were watered with Hoagland solution (11 mM NO3, 1 mM NH4) with a dripping system (1 min every 12h for the first three weeks when plants were 7-8 nodes long and 2 min every 8h afterwards). Salt stress was induced three weeks after planting and NaCl concentration was increased every 2 days (50 mM, 75 mM,100 mM,125 mM, 150 mM, 175 mM, 200 mM). Phenotype was monitored after 8 weeks after planting. 12 plants per cultivar per treatment were used.

### Ion measurement

After phenotyping, roots were rinsed in distilled water to eliminate peripheral ions originating from the vermiculite. Leaves from the fourth and fifth nodes from the apex were selected for ion measurement. Following desiccation, dry tissues were finely ground using a hammer mill with a 1 mm sieve, as described in (Nguyen et al., 2013). The twelve biological replicates obtained from phenotyping were combined in pairs, resulting in 6 final replicates per cultivar and condition. 40 ± 3 mg of tissue powder was used except for stolon node roots of which 15 mg was used. Powders were ashed at 575°C for 5 h. Ashed samples were dissolved by shaking for 30 min in 1 ml 3 M formic acid at 99°C and then diluted with 9 ml Milli-Q water. The samples were shaken again at 80°C for another 30 min. A final 500x dilution was subsequently prepared by mixing 0.3 ml sample solution (0.2 ml for the leaves) with 9.8 ml MiliQ to assess the Na^+^, K^+^, Cl^−^, and Ca2^+^ content of each root and leaf sample using Ion Chromatography (IC) system 850 Professional, Metrohm (Switzerland).

### Hydroponic experiment and sampling

We used a half-concentrated Murashige and Skoog including vitamins and 0.05 mM MES (pH=5.8) as a nutrient solution which was refreshed every two days. The plants were grown in black hydroponic containers within a growth chamber under 16/8 h light/dark light cycle at 21°C, allowing roots to grow in the dark. In the combined RNA-seq/ABA quantification experiment, salt stress (125 mM) was induced after 2 weeks at ZT4 (Zeitgeber time, 4 hours in light). Roots and leaves were collected at ZT10 (time point 6h) and the following day at ZT4 (time point 24h). Three biological replicates were used and each biological replicate is composed of the root or leaves of three plants. Tissues were flash-frozen in liquid nitrogen and subsequently ground with a mortar. For the suberin quantification, salt stress (125mM) was added to the media of two weeks old plants and roots were transferred to the fixation buffer after one week of salt stress.

### RNA extraction and RNA-seq analysis

RNA extraction and raw data analysis are available in methods S1. Differentially expressed genes (DEGs) were selected with an adjusted p-value < 0.01. The potato orthologs of the *Arabidopsis* genes were found by performing a blastp analysis using the potato protein sequence (v6.1.working_models.pep.fa), available at the SpudDB webpage, (Pham et al., 2020) as query. We used a restrictive cut-off of E-value <E-05. The potato/Arabidopsis orthologs table is available as table S1, annotations are those of TAIR10. The Gene Set Enrichment Analysis (GSEA) test of overrepresentation was performed as described in (Subramanian et al., 2005; Nicolas et al., 2022). The different GSEA lists are available in the table S2 (Gonzali et al., 2006; Baena-González et al., 2007; Osuna et al., 2007; Dinneny et al., 2008; Gifford et al., 2008; Sulpice et al., 2009; Iyer-Pascuzzi et al., 2011; Kinoshita et al., 2012; Lan et al., 2012; Li and Lan, 2015; Huang et al., 2017; Tarancón et al., 2017; Brault et al., 2019; Rymen et al., 2019; Leal et al., 2022; Nicolas et al., 2022; Lamers et al., 2023). The Gene Ontology analysis was performed using the R package topGO (p-value<0.05; Alexa and Rahnenfuhrer, 2023). GO terms of each gene come from the Arabidopsis orthologs. The transcription factor list used for the transcription factor enrichment comes from the PlantTFDB v5.0 (*Arabidopsis*) and table S1 was used to identify the orthologs in potato. We performed a hypergeometric test to determine if the enrichment is statistically significant and selected with a p-value<0.05. Arabidopsis orthologs of the potato DEGs were determined using table S1. The RNA-seq data are available in the EMBL Biostudies database available at: ebi.ac.uk/biostudies/arrayexpress/studies/E-MTAB-14181?key=42aea865-a2a9-4eec-ae78-5759443419cd. Table of the normalized count values is available as table S3.

### ABA quantification

The whole root system was quickly sampled and flash-frozen in liquid nitrogen. Leaves from the fourth and fifth nodes from the apex were selected and flash-frozen. Five biological replicates were used to quantify ABA content in root and leaves and each replicate is a pool from three plants. Samples were ground in liquid nitrogen. About 17mg of root and 30mg of leaves were used for this quantification. ABA measurement was performed as described before Li et al. (2023) and in method S2.

### Suberin deposition quantification

Fixation, clearing and staining were done according to Ursache et al. (Ursache et al., 2018). Detailed protocols of staining on mounting are available in method S3. At least 10 z-stacks images (10 focal plans) were captured for each image. Images of three different roots from each plant were taken. Image processing was performed using Fiji (Schindelin et al., 2012). Z-stack images were merged with the “sum slices” function. The root area was outlined and the mean intensity was measured. Background intensity outside the root region was also measured for quantification correction following the formula: Intensity = (Area * Mean) – ((AreaBackground * MeanBackground) * (Area / AreaBackground)) / Area. Signal intensity of five plants per cultivar and conditions were measured. Value of each biological replicate represents the mean value obtained from three roots. Five biological replicates were used for the Auramine 0 quantification and four for Yellow 088.

### NaCl-induced root K^+^ flux analysis using MIFE

Quantification of the K+ flux is described in method S4.

## Results

### Innovator is the most resilient cultivar but displays significant water-loss in response to salt

Previous studies indicated Mozart as salt-sensitive while Desirée and Innovator were identified as more resilient (Jaarsma et al., 2013; Ahmed et al., 2020). To deeper investigate the mechanisms of salt stress resilience in potato we studied these contrasting cultivars’ phenotype under a gradual salt stress in the greenhouse. After eight weeks, all cultivars showed reduced shoot length, node number, shoot branch number in salt stress condition (Figs. 1a, S1a,b,c), along with a reduction in leaf and main shoot fresh weight (FW) and leaf Normalized Difference Vegetative Index (NDVI; Figs. 1b, S1c). Among the three cultivars, Innovator was the least affected by the salt stress as illustrated by its lower reduction of leaf FW and branch number (Figs. 1b, S1b). Belowground traits showed more variation. The basal roots’ FW remained unaffected but stolon node roots’ FW was significantly reduced in Innovator and Mozart (Figs. 1c, S1d). Number of primary stolons, emerging from the main shoot, and number of secondary stolons, emerging form the primary stolons, decreased in Innovator and Mozart salt treated plants, resulting in a decreased stolon’s FW (Fig. 1d,e, S1e). Finally, Desirée and Innovator plants had a weak increase in tuber numbers under salt stress which was linked to a significant yield reduction in Desirée suggesting that the plants intend to produce more tubers to compensate the decrease of tuber’s starch accumulation (Fig. 1f, Fig. S1f).

**Fig. 1.**
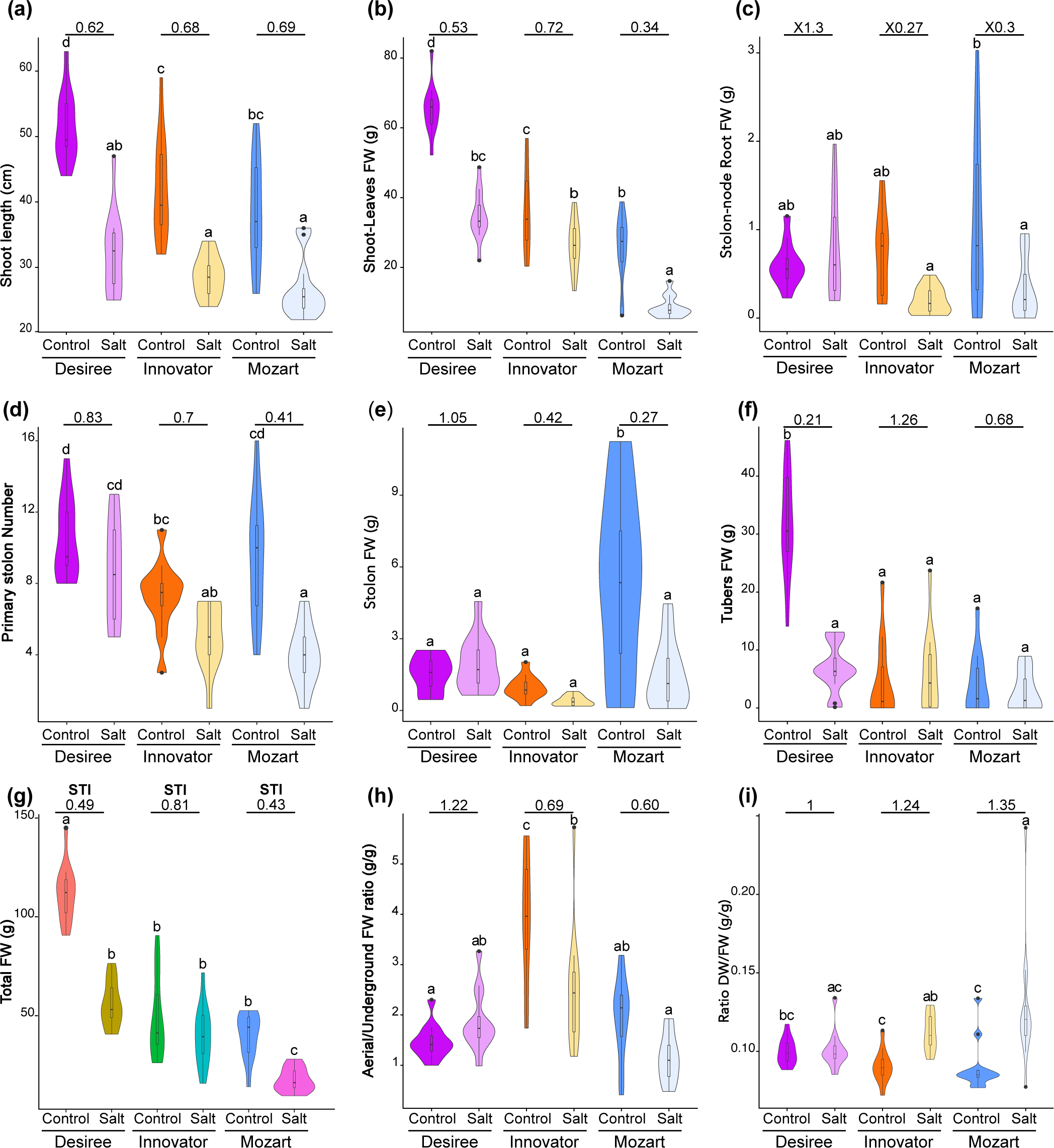
Phenotype of Desirée, Innovator and Mozart potato plants in salt stress conditions. Plants were grown in the greenhouse and phenotype monitored after 8 weeks. Salt stress was induced three weeks after planting and NaCl concentration was increased every 2 days (50 mM, 75 mM,100 mM,125 mM, 150 mM, 175 mM, 200 mM). (a-i) Violin plots representing diverse above and below ground phenotypes (n=12). (a) length of the shoot. (b,c,e,f,g) Fresh weight (FW) of the shoot and leaves (b), stolon-node root (c), stolon (e), tubers (f) and total (salt-tolerance index, STI; g). (d) number of primary stolons emerging from the shoot below ground. (h) ratio of aerial part (shoot and leaves) FW/below ground FW (including basal and stolon-node roots, stolons and tubers). (i) ratio of dry weight (DW)/(FW) as indicator of the water retention. Value on top of each Control-Salt plot represents the salt/Control ratio. The letters designate significant differences among means (one-way ANOVA plus Tukey’s HSD).

Next, we calculated the salt-tolerance index (STI), the ratio of total fresh weight between salt and control conditions, in order to evaluate how salt globally affects the cultivars. The data showed that Innovator has the highest STI followed by Desirée suggesting they are more tolerant than Mozart (Fig. 1g). We also calculated the ratio of fresh weight between aerial and underground parts to assess the effect of salt stress to biomass partitioning responses. In control conditions, Innovator plants had the highest aerial/underground ratio (Fig. 1h), suggesting a greater allocation of energy aboveground in this cultivar compared to Desirée which had the lowest. In salt-treated plants, this ratio decreased in Innovator and Mozart plants, mostly as a consequence of their more conserved tuber yield in salt stress conditions. Likewise, Innovator and Mozart showed an increased dry weight DW/FW ratio of the entire plant suggesting that they displayed a lower water retention under salt stress than Desirée (Fig. 1i).

Ultimately, we performed a correlation analysis between the different traits. Aside the expected positive correlation between shoot and leaves traits, results showed several positive correlations between the different stolon traits, especially in Innovator and Mozart (Fig. S1g-i). In particular, stolon weight and stolon root weight were correlated which suggested their growth is highly linked. However, tuber number and weight were not correlated with any other subterranean traits, except in Innovator where it is weakly correlated with basal root weight. Nevertheless, these tuber traits were correlated with leaf weight in Desirée and Innovator. This suggested that tuber production, as expected, depends on sugar production. The association with any other underground part is however more complex.

In conclusion, this phenotypic analysis highlights the differential responses and plasticity of the three cultivars to salt stress. Desirée, despite having the second-highest STI, experienced significant reduction in leaf weight and tuber yield but maintained overall a good water content. Conversely, Innovator and Mozart, which initially have a carbon allocation turned towards the aerial part, exhibit similar yields under both control and salt stress conditions, particularly Innovator. Due to its low STI, lower tuber yield even in control conditions and its stronger water loss, Mozart does not represent an interesting cultivar. On the other hand, Innovator appears to be the most resilient cultivar with the highest STI and its unchanged tuber yield in salt stress. It is likely the most interesting cultivar from the agricultural point of view although this resilience is associated with a significant water-loss under salt stress.

### Innovator has the most divergent acclimation transcriptomic response

To understand in more detail the molecular mechanism of the salt stress response in the adventitious roots of the three cultivars, we conducted an RNA-seq transcriptomic analysis. We compared the transcriptomes of the basal roots of the three cultivars grown in control or salt stress conditions (125mM), 6h and 24h after stress induction. Principal component analysis (PCA) showed that the first principal component (PC1) separated the control and treated roots of Mozart and Desirée plants after 6h in contrast to Innovator which also displayed fewer differentially expressed genes (DEGs; Fig. 2a,b; table S4). After 24h, PC1 marked a clear salt stress response in the three cultivars, including Innovator which exhibited the highest DEG number (Fig. 2a,b; table S4). This temporal development of transcriptome patterns hence suggested a dynamic acclimation process to salt stress across the cultivars.

**Fig. 2.**
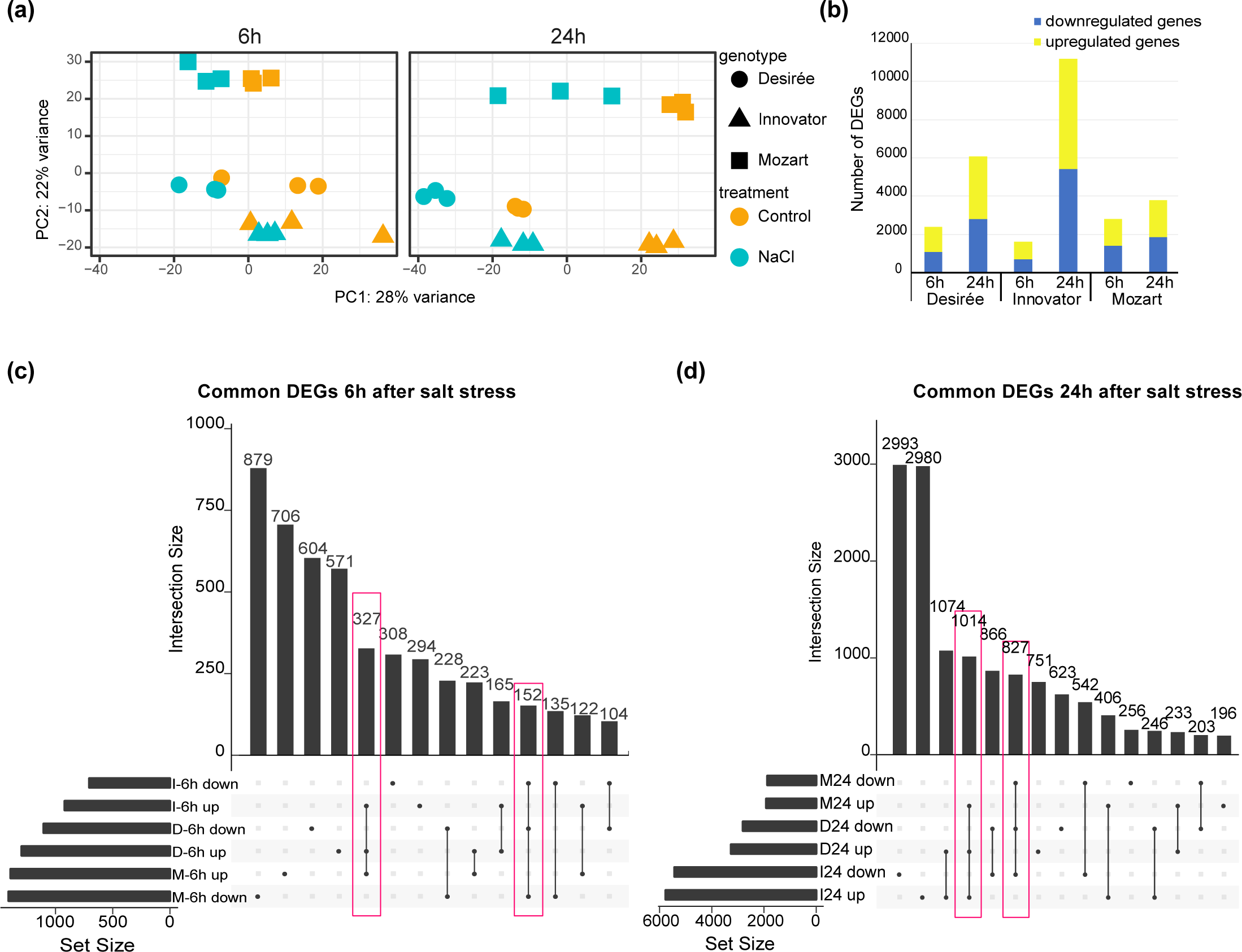
Overview of the transcriptomic response observed in the three cultivars after salt stress (125mM). (a) Principal Component Analysis (PCA) of the salt stress response of the three cultivars (Desirée, Innovator, Mozart) at 6h and 24h. (b) Number of up-regulated and down-regulated Differentially Expressed Genes (DEGs, adj. p-values<0.01) at 6h and 24h. (c-d) UpSet graph (Lex et al., 2014) showing common DEGs between different cultivars and treatment at 6h (c) and 24h (d).

The comparison of the DEGs between the cultivars showed that after 6h, 327 upregulated genes were shared between the three cultivars, while only 152 downregulated genes were shared (Fig. 2c). In order to identify the genetic pathways involved in salt stress responses we performed a Gene Ontology (GO) enrichment analysis and a Gene Set Enrichment Analysis (GSEA) which is a statistical approach allowing the identification of over-represented gene sets among the up- and downregulated DEG (Table S2; Subramanian et al., 2005). The shared upregulated genes expectedly showed an enrichment for several stress-related terms including water deprivation, salt stress, cold, wounding and osmotic stresses as well as K^+^ homeostasis (tables 1, S5). Consistently, GSEA revealed that down- and upregulated gene sets previously associated with Na^+^ stress response in Arabidopsis were accordingly enriched in our datasets, emphasizing the conservation of salt stress response in plants (Figs. 3a, S2; Dinneny et al., 2008; Lamers et al., 2023). Likewise, sets of genes related to different stresses were also found enriched, underlying the existence of a core gene set in common between the different abiotic stress (Figs. 3a, S2; Dinneny et al., 2008; Iyer-Pascuzzi et al., 2011; Lan et al., 2012; Rymen et al., 2019). Interestingly, Innovator displayed a milder stress response after 6h suggesting a distinct mechanism.

**Fig. 3.**
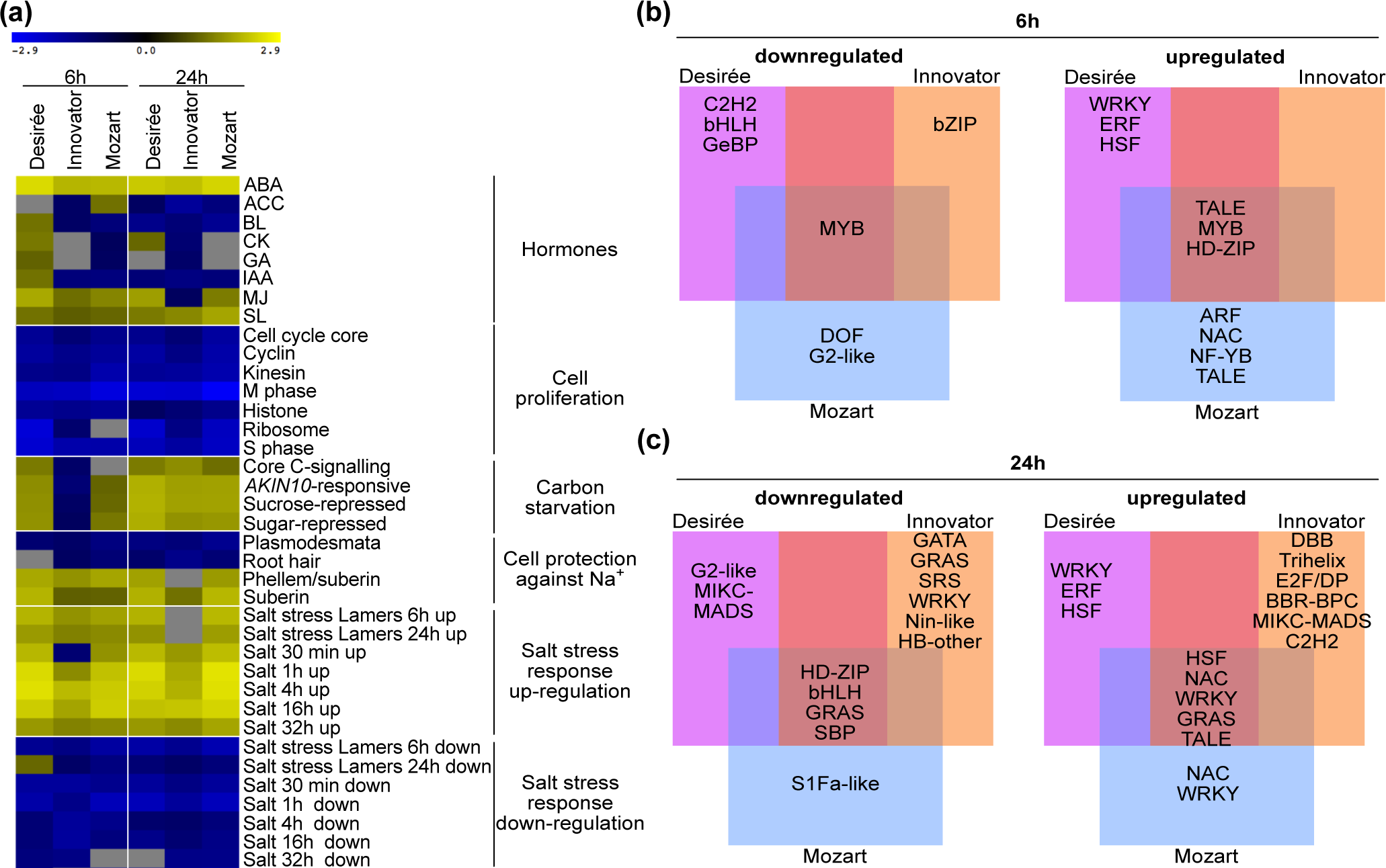
Transcriptomic changes in response to salt stress in Desirée, Innovator and Mozart. (a) Heatmap of the Gene Set Enrichment Analysis (GSEA) displaying the normalized enrichment score (NES). GSEA is a statistical approach allowing the identification of over-represented gene sets among the up- and downregulated DEG. Gene sets come from the literature, mostly in *Arabidospsis*, and available in table S6. Positive NES values (yellow) indicate gene sets which are overrepresented among induced genes and Negative NES values (blue) indicate those which overrepresented among the repressed genes. Null values indicate gene sets not significantly overrepresented. (b-c) Venn diagrams showing the transcription factor families overrepresented (p-value <0.05 in a hypergeometric test) among the up-regulated and down-regulated DEGs in the three cultivars at 6h (b) and 24h (c).

Additionally, both GO and GSEA analyses highlighted the importance of ABA and jasmonic acid signalling in salt stress response after 6h (tables 1, S5, Fig. 3a). GSEA also indicated a homogenous activation of the Strigolactones (SL) signalling pathways (Fig. 3a). On the other hand, GO and GSEA gene set related to cell division and growth processes were logically downregulated (Fig. 3a, tables 1, S5). Likewise, plasmodesmata and root hair development gene sets were inactivated (Huang et al., 2017; Brault et al., 2019) while suberin deposition genes (Leal et al., 2022) were activated indicating that the cellular acclimation aimed to limit Na^+^ absorption and diffusion in the roots (Fig. 3a). Interestingly, gene sets related to carbon starvation (Gonzali et al., 2006; Baena-González et al., 2007; Osuna et al., 2007; Sulpice et al., 2009; Tarancón et al., 2017) were found activated in Desirée and Mozart after 6h which is consistent with a lower sugar loading in the roots due to the growth arrest (Fig. 3a). Although cell proliferation gene sets are downregulated in Innovator, the carbon starvation gene sets are surprisingly also downregulated in this cultivar after 6h emphasizing the singular response dynamic of this cultivar.

After 24 hours exposure to salt stress, 1014 upregulated and 827 downregulated genes were found in common between the three cultivars (Fig. 2d). A continued activation of stress-related, suberin deposition and ABA signalling genes was consistently observed (tables 1, S5, Fig. 3a). Carbon starvation response was activated suggesting that it was previously delayed. GO terms analysis showed that lateral root growth, water and symplastic transport were downregulated which is consistent with the persistent inactivation of plasmodesmata and root hair growth genes sets observed (tables 1, S5, Fig. 3a). Moreover, downregulation of ethylene and brassinosteroid (BR) signalling pathways was evident across cultivars (Fig. 3a), while cytokinin (CK), gibberellins (GA) and methyl jasmonate (MJ) were uniquely downregulated in Innovator which further highlights its distinct response profile (Fig. 3a).

Suberin and K^+^ efflux measurements suggested potential differences in Na^+^ influx and Na^+^/K^+^ homeostasis between the cultivars. To identify which channel/transporter genes are involved in salt acclimation in potato, we investigated the expression pattern of the different genes identified. Results showed that SKOR/GORK K^+^ efflux channel and AKT1 K^+^ influx coding genes were upregulated during the stress response of the cultivars suggesting they play a role in the shared salt response (Fig. S3). Interestingly, another SKOR/GORK gene was found more expressed in Innovator and down-regulated in salt stress conditions, which lead to the assumption that it could be involved in the initial strong K^+^ leakage in Innovator. Several transporters from different classes like AKT1, HAK7, NHX5, NHX6 and KEA3 and KEA4 were specifically upregulated in Innovator after 24h indicating they may play an important role in the salt acclimation of this genotype.

Altogether, our transcriptomics data showed that the salt stress response exhibits a shared core mechanism between *Arabidopsis* and potato, and also has a part in common with other abiotic stresses. Innovator shows a unique temporal pattern coupled with a differential activation of stress-related pathways and channel/transporter genes which point to a distinct resilience strategy in this cultivar.

### The different salt stress response phases specifically involve different TF families

Because of their important roles in regulating gene networks, we examined transcription factor (TF) family’s enrichment in response to salt stress (Fig. 3b). We found distinct patterns emerging over time and across different cultivars. After 6h, the *TALE* (*Three Amino acid Loop Extension*), the *MYB* (*MYeloBlastosis*), and the *HD-ZIP* (*Homeodomain-leucine Zipper*) family were enriched in the shared upregulated genes suggesting a conserved early activation of these TFs in response to the stressor (Fig. 3b, table S6). In addition, the stress-related TF families *WRKY*, *HSF* (Trimerization of heat shock transcription factor) and *ERF* (*Ethylene Responsive Factor*) were enriched in Desirée suggesting an early activation of stress-adaptation pathways in this cultivar. After 24h, the TF landscape showed a more intricate picture. HD-ZIPs were subsequently downregulated indicating a specific role in the early stage of stress response (Fig. 3c, table S6). While TALE continued to be upregulated, additional stress-related TF families like *HSF* and *WRKY* were also enriched, along with *GRAS* and *NAC* (*NAM*, *ATAF1/2*, and *CUC2*) which are usually involved in growth and development. This indicated a broader and more intensified transcriptomic response in roots, likely corresponding to the growth recovery phase. Interestingly, Innovator displayed unique TF enrichment patterns, confirming its major reprograming and recovery phase at this time point.

Altogether these results reveal a nuanced and cultivar-specific regulation of TFs within the first 24h of salt stress in the roots. The stress response first includes the recruitment of specific TF families like MYB and HD-ZIP at 6h of stress, followed by HSF, WRKY, NAC and GRAS at 24h, while TALE TF played a continuous role.

### Tubers do not accumulate sodium under salt stress

In addition to the traditional shoot, leaf and root tissues, potato plants develop different underground organs like stolons, stolon node roots and tubers all in contact with salt in the soil. To determine whether these organs differentially accumulate toxic ions under salt stress, we measured the ion contents of the different organs after the plant phenotyping (Fig. 1). As expected, sodium (Na^+^) and chloride (Cl^-^) contents were significantly increased in the two kinds of root as well as in the stolon of the three cultivars (Fig. 4a-c). Na^+^ concentration also rose in Mozart and Innovator aerial part but remained unchanged in Desirée (Fig. 4d). Surprisingly, Na^+^ content in tubers showed no significant difference between cultivars while Cl-concentration was consistently higher, suggesting an exclusion of Na^+^ from tubers in salt conditions (Fig. 4e). Interestingly, potassium (K^+^) content also increased in tubers resulting in a mildly decreased Na^+^/K^+^ ratio (Fig. 4e), while it decreased in basal roots and in some aspects in stolon node roots, leading to an increased Na^+^/K^+^ ratio (Fig. 4a,b). This ratio increased in Innovator and Mozart leaves, due to their higher Na^+^ and lower K^+^ content (Fig. 4d). Calcium (Ca2^+^), phosphate (PO_4_2^-^) and sulphate (SO_4_2^-^) concentrations remained unchanged between control and salt treatments, suggesting that they do not play any role in long term salt stress adaptation (Fig. S4a-e). However, magnesium content was significantly decreased in the aerial part (shoot and leaves) of Mozart plants (Fig. S4d) while it increased in stolons suggesting that it might have a role in salt stress adaptation in this cultivar.

**Fig. 4.**
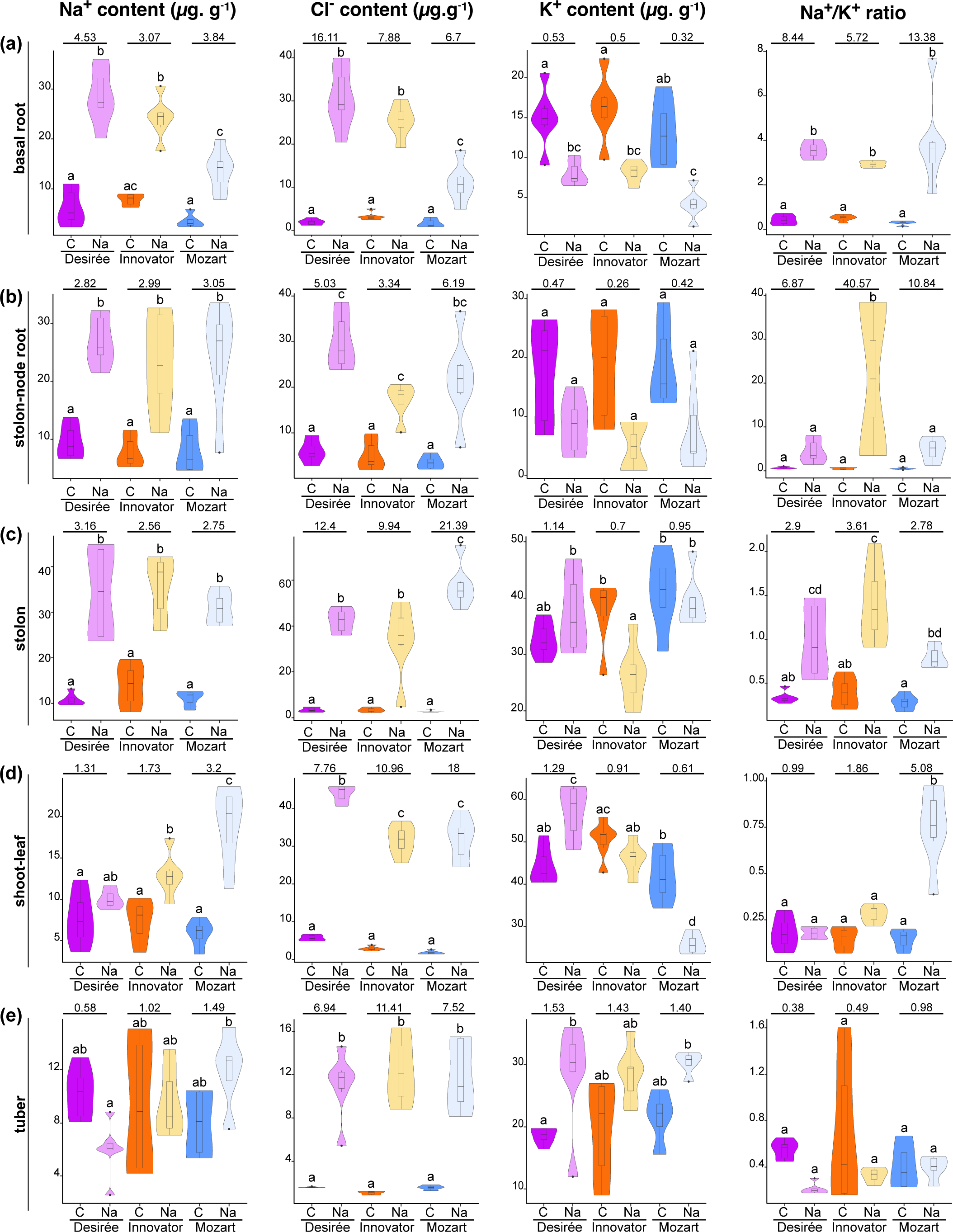
Na+, Cl- and K+ content (µg.g-1) in Desirée, Innovator and Mozart after salt stress. Na+, Cl- and K+ contents in basal and stolon-node roots (a and b respectively), stolon (c), shoot land leaves tissues (d) and tubers (e). Sample size is constituted by 6 replicates, each of them forms by a pool of two plants. Plants used are those of the phenotyping (Fig. 1) and grown in the greenhouse in control conditions (C) or with an increased NaCl stress (Na) up to 200mM. The letters designate significant differences among means (one-way ANOVA plus Tukey’s HSD).

These results indicate varied strategies for Na^+^ accumulation among cultivars. Aboveground, Na^+^ likely accumulates in the leaves/stems of Innovator and Mozart. Belowground, root types and stolons accumulated Na^+^ which is actively excluded from the tubers that hence do not represent an additional organ for Na^+^ sequestration.

### Innovator initially exhibits a lower suberin and lignin deposition level and a higher K**^+^** leakage but adequality acclimates to salt stress

Suberin and lignin can be deposited in the cell wall of specific root cells and acts as a barrier preventing radial ion transport to the stele upon stresses (Ranathunge and Schreiber, 2011; Barberon et al., 2016; Doblas et al., 2017). In salt-stressed roots, Na^+^ content increased while K^+^ decreased, in particular in basal roots (Fig. 4a,b). GSEA showed that suberin gene set is activated in the three cultivars but that the intensity of the activation varies suggesting variation of suberin deposition (Fig. 3a). We hence confirm that suberin and lignin deposition occurs in salt-stressed potato roots. We stained basal roots with Nile Red and Yellow 88, specifically staining suberin; Auramine O that stains lignin, suberin and cutin; and Acridine orange which stains lignin, cutin, cellulose, DNA and the polyphenolic domain of suberin (Briggs and Morris, 2008; Kováčik et al., 2014; Houtman et al., 2016; Ursache et al., 2018). We could not detect any staining with Nile red (Fig. S5a) while conversely, Yellow 88, Auramine O and acridine orange stained the differentiated zone of the root (Fig. S5a). Orthogonal view of basal root stained with Auramine O indicated that the suberin and lignin deposition took place in the exodermis and in the Casparian strips (Fig. S5b). We next quantified the suberin and lignin deposition of the roots of the three cultivars one week after salt stress induction using yellow 88 and Auramine O. In control conditions, Mozart showed a higher suberin and lignin deposition compared to the other two cultivars suggesting that it may be more protected against salt absorption (Fig. 5a,b). However, after one week of salt treatment, these depositions were unchanged in Mozart’s roots while Innovator roots adjusted their deposition levels under salt stress, eventually reaching levels comparable to those observed in Mozart roots suggesting that Innovator roots can adapt their suberin and lignin levels upon salt stress (Fig. 5a,b). As the increased deposition was only significant when roots were stained with Auramine O, it is likely that lignin deposition is important for the root acclimation to salt stress in Innovator. Similarly, Desirée root acclimate to salt stress by increasing suberin and lignin deposition but this accumulation does not reach the level of Mozart and innovator under salt stress.

**Fig. 5.**
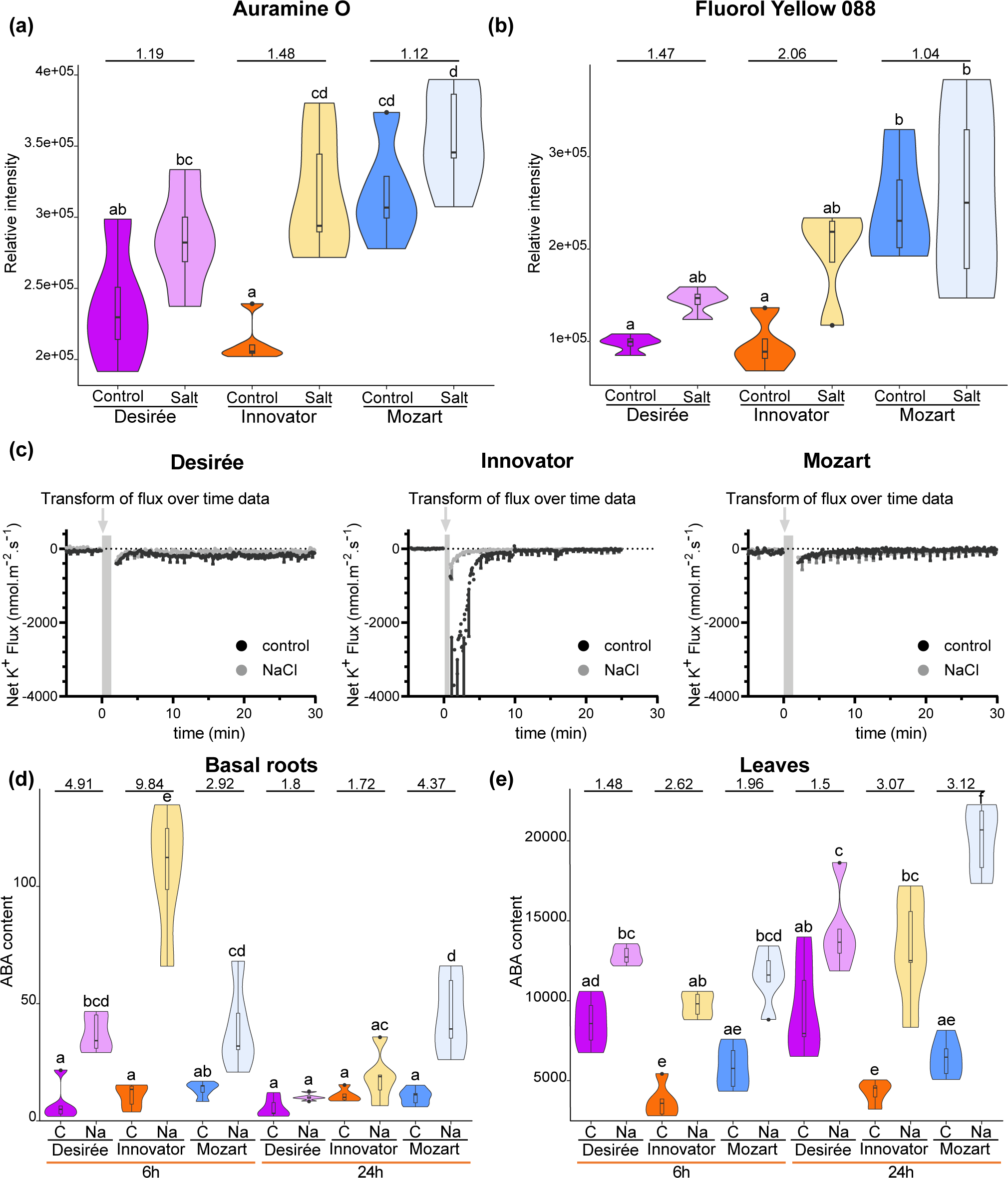
Plant response to salt stress. (a-b) Quantification of suberin and lignin deposition in the basal root of the three cultivars. 20-day old plants were grown in *in vitro* in control or salt stress conditions (125mM) for one week. Auramine O (lignin and suberin) or Fluorol Yellow 088 (suberin) staining reagents were used (n=5, each replicate represents the mean value of 3 roots). (c) K+ flux in basal roots measured in control plants or plants previously stressed (75mM) for 7 days. Five biological replicates per treatment and genotype were used. Data represent the mean, error bars the standard deviation. (d-e) Violin plots depicting ABA contents in basal roots and leaves of the three cultivars in response to salt stress after 6h and 24h (n = 5 pools of 3 plants). Plants were grown in hydroponic conditions in control conditions (C) or in salt stress conditions (Na). Values are normalized by fresh weight of the samples. The letters designate significant differences among means (one-way ANOVA plus Tukey’s HSD).

These results suggested that Innovator and Desirée are conceivably less protected against sudden Na^+^ stress than Mozart which hence could lead to a rapid Na^+^ influx and osmotic stress. To explore whether this difference of suberin deposition can also influence the Na-induced K^+^ efflux, we used a non-invasive microelectrode ion flux (MIFE) system to measure K^+^ efflux in response to NaCl addition (75mM). In control conditions, Innovator roots showed a significant K^+^ efflux which was not observed in Desirée or Mozart (Fig. 5c). However, after 14 days of salt stress acclimation, Innovator roots no longer exhibited this net K^+^ efflux from root cells, suggesting that Innovator can acclimate to NaCl by decreasing K^+^ efflux under salt stress.

In conclusion, root suberin and lignin depositions vary between the roots of the cultivars under normal conditions. Mozart displays higher levels in the exodermis and Casparian strips even before salt stress induction. Innovator demonstrated a remarkable ability to acclimate to salt stress. It promptly increases the suberin content from the initial lowest content to a quantity similar to that of Mozart and limits its root K^+^ loss in response to salt.

### ABA content indicates the degree of salt stress response and salt acclimation in the three cultivars

ABA is a well-known hormone mediating stress responses in plants (Trivedi et al., 2016). GO Analysis and GSEA indicated an activation of ABA signalling in roots after salt stress (Table 1, Fig. 3a) To confirm its role in salt stress responses in potato, we measured the ABA content in roots after 6h and 24h. The results confirmed our hypothesis that ABA concentration is increased in salt-treated plants compared to the control (Fig. 5d). However, the increased concentration of ABA was stronger in Innovator roots at 6h compared to the other cultivars. After 24h of treatment, the level of ABA was equivalent to the control in Desirée and Innovator roots. However, in Mozart the ABA content remained high suggesting that Mozart roots were still in a stress phase after 24h of stress.

**Table 1.** GO analysis of salt stress response. Gene Ontology (GO) analysis of salt stress response illustrating the ten most enriched biological process GO terms for DEG shared among three cultivars. The enriched GO terms for down-regulated and up-regulated genes are presented separately.

|  | common downregulated | significant | p-value | common upregulated | significant | p-value |
| --- | --- | --- | --- | --- | --- | --- |
| 6h | DNA replication initiation | 8 | 6.10E-11 | response to water deprivation | 81 | 1.40E-13 |
|  | double-strand break repair via break-induced replication | 4 | 2.60E-06 | response to abscisic acid | 85 | 2.70E-11 |
|  | anatomical structure arrangement | 11 | 6.20E-06 | response to salt stress | 51 | 1.00E-09 |
|  | nuclear DNA replication | 4 | 7.70E-06 | positive regulation of syringal lignin biosynthetic process | 8 | 1.30E-08 |
|  | organic acid metabolic process | 47 | 0.00018 | response to oxidative stress | 40 | 1.20E-07 |
|  | plant organ morphogenesis | 18 | 0.00083 | defense response to fungus | 47 | 7.60E-07 |
|  | response to desiccation | 4 | 0.00116 | positive regulation of seed germination | 9 | 2.10E-06 |
|  | intracellular iron ion homeostasis | 4 | 0.00135 | response to wounding | 52 | 2.60E-06 |
|  | DNA unwinding involved in DNA replication | 3 | 0.00152 | jasmonic acid and ethylene-dependent systemic resistance | 6 | 7.00E-06 |
|  | response to inorganic substance | 43 | 0.00261 | response to cold | 40 | 1.30E-05 |
| 24h | double-strand break repair via break-induced replication | 6 | 1.20E-07 | response to water deprivation | 136 | 6.10E-15 |
|  | DNA replication initiation | 8 | 1.40E-07 | response to abscisic acid | 134 | 2.20E-10 |
|  | macromolecule biosynthetic process | 194 | 4.70E-07 | response to wounding | 90 | 5.10E-10 |
|  | cell wall biogenesis | 56 | 5.90E-07 | cellular catabolic process | 176 | 7.10E-10 |
|  | plant-type cell wall biogenesis | 35 | 1.70E-05 | organonitrogen compound catabolic process | 113 | 1.30E-09 |
|  | mitotic cell cycle process | 26 | 4.70E-05 | cellular response to hypoxia | 41 | 1.60E-09 |
|  | unidimensional cell growth | 34 | 6.30E-05 | response to oxidative stress | 84 | 3.70E-09 |
|  | microtubule cytoskeleton organization | 17 | 9.00E-05 | response to salt stress | 69 | 1.10E-08 |
|  | plant-type cell wall organization or biogenesis | 63 | 9.70E-05 | leaf senescence | 43 | 1.20E-07 |
|  | regulation of actin filament depolymerization | 4 | 0.00013 | defense response to bacterium | 92 | 1.60E-07 |

After induction of salt stress in the roots, ABA can be transported and/or synthesised in other part of the plant, for example in leaves (Trivedi et al., 2016). Thus, to study the ABA accumulation pattern in potato cultivars, we also quantified its content in leaves. The results showed that ABA content was increased in the three cultivars after 6h (Fig. 5e). However, the ABA content in the leaves remained significantly higher in the three cultivars compared to the control after 24h in contrast to the roots. The difference between treated and control plants was actually stronger in Innovator and Mozart suggesting that these two cultivars induced a stronger ABA-related stress response which is consistent with their higher sodium content in the aerial parts (Fig. 4d).

In conclusion, the higher ABA content in the roots in Innovator is consistent with a stronger initial stress response in this cultivar and can be associated with its initially lower suberin deposition in root cell wall and K^+^ leakage. The higher ABA content in leaves in Innovator and Mozart is compatible with the higher Na^+^ storage in the leaves of these two cultivars and suggest a stronger stress response to salt.

## Discussion

This study combined plant phenotyping, physiological, transcriptomic, metabolomics analysis and confocal imaging to explore how three contrasting cultivars respond and acclimate to salt stress. In addition to the classical phenotyping of the leaves and roots, we included the stolons, stolon node roots and tubers, which are usually excluded from such studies in potato. These results showed than cultivars differently respond to salt stress, highlighting different ability of acclimation and resilience.

### Innovator and Desirée, two distinct responses to salt stress

Determining the most resilient genotype depends on the traits of interest. Potato agronomic utility is based on tubers production. Consequently, tuber yield is a pivotal criterion for identifying resilient potato cultivars. Under salinity stress, Innovator was the most resilient cultivar with minimal tuber yield changes and the highest STI. However, its low water retention could be disadvantageous under field conditions.

Conversely, Desirée, thought showing reduced tuber yield, demonstrated significant resilience with the second-highest STI and the best water retention, which confirms previous findings (Jaarsma et al., 2013). Mozart was the most sensitive genotype and shows the highest Na^+^/K^+^ ratio in leaves. The higher Na^+^ concentration and ABA content in Innovator and Mozart leaves after 24h (Fig. 4d) could results from higher water loss due to a higher transpiration stream and hence Na^+^ transport to this organ (Asch et al., 2000; Jaarsma et al., 2013).

Suberin and lignin coats, crucial for Na^+^ exclusion in many species (Ranathunge and Schreiber, 2011; Barberon et al., 2016; Karlova et al., 2021; Li et al., 2023), are less deposited in Desirée and Innovator. It potentially leads to a higher root hydraulic conductivity and hence greater absorption of Na^+^ and Cl^-^ ions within the first hours, generating more stress. A SKOR/GORK channel is more expressed in Innovator and could likely be involved in the strong net K^+^ efflux observed in Innovator’s control roots (Fig. S3). Other identified channels and transporter genes in Innovator are interesting candidates for studying salt stress acclimation in potato.

In the meanwhile, ABA accumulation may contribute to the compensatory suberin deposition observed at 24h in Innovator and Desirée and hence to the Na^+^ influx limitation (Barberon et al., 2016; Shukla et al., 2021). In terms of salt-stress resilience, this result indicates that the initial suberin content under control conditions and the associated high root K^+^ efflux or loss is less critical than the roots’ ability to adequately acclimate to the stress.

### Deciphering the early steps of salt stress response in potato

Innovator’s ABA and suberin levels, along with K^+^ efflux suggest a longer root quiescent phase in this cultivar which could explain its different transcriptomic pattern. After 24h, this delayed salt response might lead to a prompt and substantial transcriptome reprogramming of almost a third of the total gene number and could explain why sugar starvation and wounding gene sets are not activated after 6h in Innovator. Nonetheless GO and GSEA identified conserved pathways involved in salt-stress response in potato. Besides the expected ABA response, a homogenous activation of strigolactone (SL) pathway was identified. SLs, produced by roots, modulate the root system architecture by inhibiting adventitious and lateral root formations (Rasmussen et al., 2013). Additionally, they can relieve the negative effects of adverse environmental conditions like drought, heat and salt stress in *Arabidopsis* (Ha et al., 2014; Salvi et al., 2021). MJ, known to increase under abiotic stress conditions and likely to collaborates with ABA, is logically activated across all cultivars after 6h (Salvi et al., 2021). Moreover, MJ can reduce Na^+^/K^+^ ratio in roots and leaves in maize (Rehman et al., 2023) which is consistent with its early activation here. Unexpectedly the stress-related ethylene pathway (Achard et al., 2006) was inactivated after 24h. However, the loss of function of ethylene receptors ETHYLENE RESPONSE 1 (ETR1) and ETHYLENE INSENSITIVE 4 (EIN4), generate enhanced salt tolerance and ethylene also has a negative effect on suberin deposition and HAK5 expression (Peng et al., 2014; Van Zelm et al., 2020). Consequently, local inactivation of the ethylene pathway could take part of the root recovery phase in potato. The diverse BR, CK, and GA responses to salt stress among cultivars, suggest they have a nuanced role in the stress response. Many gene sets activated in Arabidopsis under salt stress are similarly induced in potato and highlights that salt stress response is well conserved. Furthermore, induction of phosphate and Fe starvation gene sets upon salt stress aligns with the reduce uptake of essential nutrients due to Na^+^ accumulation in roots and highlights the connection between salt stress response and nutrient uptake pathways (Parihar et al., 2015). Overall, there is a lower conservation of downregulated gene sets between *Arabidopsis* and potato which suggests they are more species-specific and likely depend on the transcriptome of the particular cultivar.

### An organised regulation of TF activity during salt stress response

Our study reveals a time-specific roles of TF families in response to salt stress. In the first hours MYBs undergo significant turnover. MYB is a large family involved in primary and secondary metabolism, cell identity, developmental processes, but also in response to biotic and abiotic stress. Several MYBs were shown to be involved in salt and osmotic stress, suberin deposition, dehydration and phosphate starvation responses and their action is often related to ABA signalling (Dubos et al., 2010; Shukla et al., 2021; Li et al., 2023). Likewise, HD-Zip TF family regulates the expression of downstream stress-related genes through ABA and their role in salt stress has already been pointed out in different species (Ariel et al., 2010; Sharif et al., 2021; Li et al., 2022). In potato, they might play a crucial role in the early stage of the salt stress response before being downregulated after 24h. TALE TFs shows consistent upregulation at both time points. A recent model states they might create a permissive chromatin platform permitting the binding of other specific TFs afterwards (Bobola and Sagerström, 2022). We can speculate that salt stress and the implied transcriptome reprogramming hence requires the preliminary action of TALE proteins. The enrichment of HSF and WRKY TF families at 24h suggests their vital role during the later acclimation phase to salt stress.

### Maintenance of a strong sugar sink belowground is a crucial process in potato under stress

Despite its high concentration in stolons, tubers did not accumulate Na^+^ The intense sugar metabolism during tuberization is likely incompatible with Na^+^ storage. Furthermore, the increased Cl- and K^+^ concentrations in the tubers strongly suggests an active Na^+^ exclusion linked to a compensatory K^+^ uptake which could involve Na^+^ and K^+^ transporters like HKT1 and HAK5 (Van Zelm et al., 2020).

Nevertheless, salt can reduce stolon and stolon node root growth and tuber yield as observed in Desirée suggesting a lower sugar sink strength belowground in those conditions. This aligns with the growth reduction and the activation of carbon starvation gene sets after salt stress observed in the GO and GSEA analyses. However, basal roots growth was not reduced, possibly due to plant adaptation during the first three weeks. Immediate salt stress induction upon tuber planting may accentuate belowground differences.

Potato tubers are located belowground, close to the roots, where salt is absorbed. Therefore, maintaining a sufficient belowground sugar sink is critical for maintaining tuber yield. Despite Desirée showing the highest yield under control conditions, tuber production decreased significantly under salt stress and became similar to Mozart and Innovator. Halophyte species like *Schrenkiella parvula* have more sugar transporter genes to compensate for decreased sugar sink (Zou et al., 2017). Therefore, an increased sugar transport belowground or enhancing starch biosynthesis in tubers may boost yield in cultivars like Desirée.

Altogether, we showed that Innovator was the most resilient cultivar due to a high degree of acclimation but displays a more severe water-loss under salt stress. Absence of Na^+^ accumulation and K^+^ increase in tubers growing under salt stress is a remarkable phenomenon that require further studies and can lead to a better understanding of salt tolerance mechanisms. Our RNA-seq analyses pointed towards different TFs and ion transporters which are interesting candidates for further analysis on the quest of breeding for salt resilience in potato.

## Acknowledgments

This work was supported by the NWO-TTW (Grant No. 16893), “Getting to the roots of salt stress resilience of potato plants”. We thank J. Busscher for the lab management at the Plant Physiology laboratory (WUR), S. Geurts and people of Unifarm for their technical assistance in the greenhouse (WUR). We are thankful to Gerard van der Linden (WUR) for useful discussions.

## Author contributions

MN performed most of the experiments and analysed the data. SEL and JBo set up and measured the suberin deposition in roots. JHV, MS, JTME and BK contributed the K^+^ flux/MIFE experiments, HvdS performed the ion measurements. FV and TdZ measured the ABA contents. DB and JBo and JBu helped with the phenotyping. DB assisted the RNAseq analysis and set up the GO analysis in potato. MN, RK and CT designed the experiments. MN and RK wrote the article, with input from CT, JHV, JB, SEL, JTME. RV, JTME, RK and CT supervised the project.

## Data availability

The data that support the findings of this study are available in the Supporting Information of this article. The sequence data that support the findings of this study are openly available in the BIOSTUDIES database (https://www.ebi.ac.uk/biostudies/) under accession no. E-MTAB-14181 ( https://www.ebi.ac.uk/biostudies/arrayexpress/studies/E-MTAB-14181?key=42aea865-a2a9-4eec-ae78-5759443419cd).

## *New Phytologist* Supporting Information

**Table S1** Potato/Arabidopsis orthologs table

**Table S2** Gene lists of the different Gene sets used for the GSEA

**Table S3** Table of the Normalized counts at 6h and 24h

**Table S4** Differentially expressed genes (DEG) lists of the three cultivars

**Table S5** Gene Ontology analysis of the salt stress response in the three cultivars

**Table S6** Transcription factors enrichment analysis

**Fig. S1.**
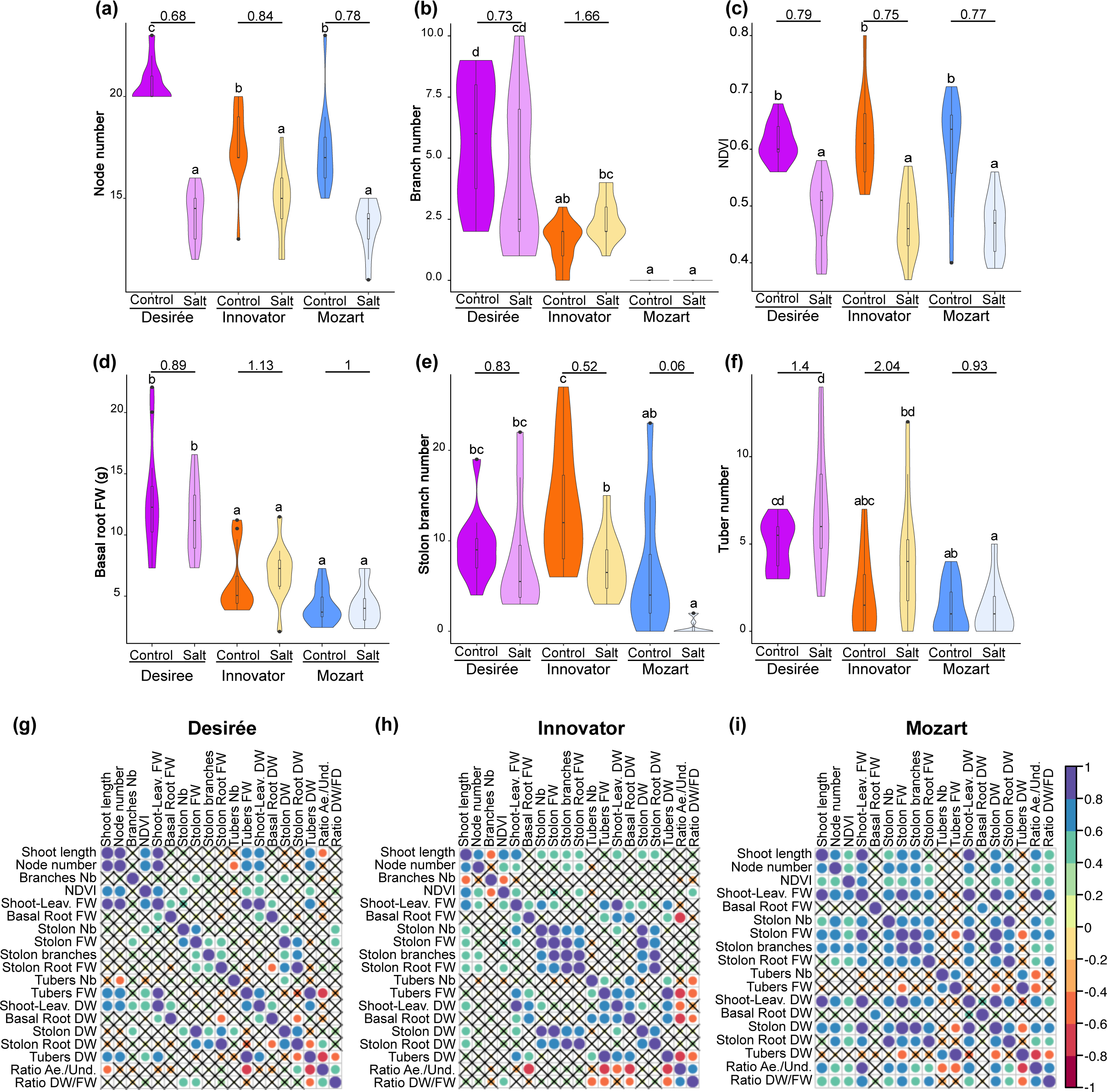
Phenotype of Desirée, Innovator and Mozart in salt stress conditions. Violin plots representing diverse above and below ground phenotypes (n=12). Nodes number (a) and branches number (b) of the aerial shoot. (c) Normalized difference vegetation index (NDVI) of the leaves. (d) FW of the basal roots. (e) Number of stolon branches emerging from the primary stolons. (f) tuber numbers collected for each plant. Value on top of each Control-Salt plot represents the salt/Control ratio. The letters designate significant differences among means (one-way ANOVA plus Tukey’s HSD). (g-i) Graphical representation of the Pearson’s correlation results. Crosses indicate the correlation which are not significant. Colours indicate the Pearson’s coefficient.

**Fig. S2.**
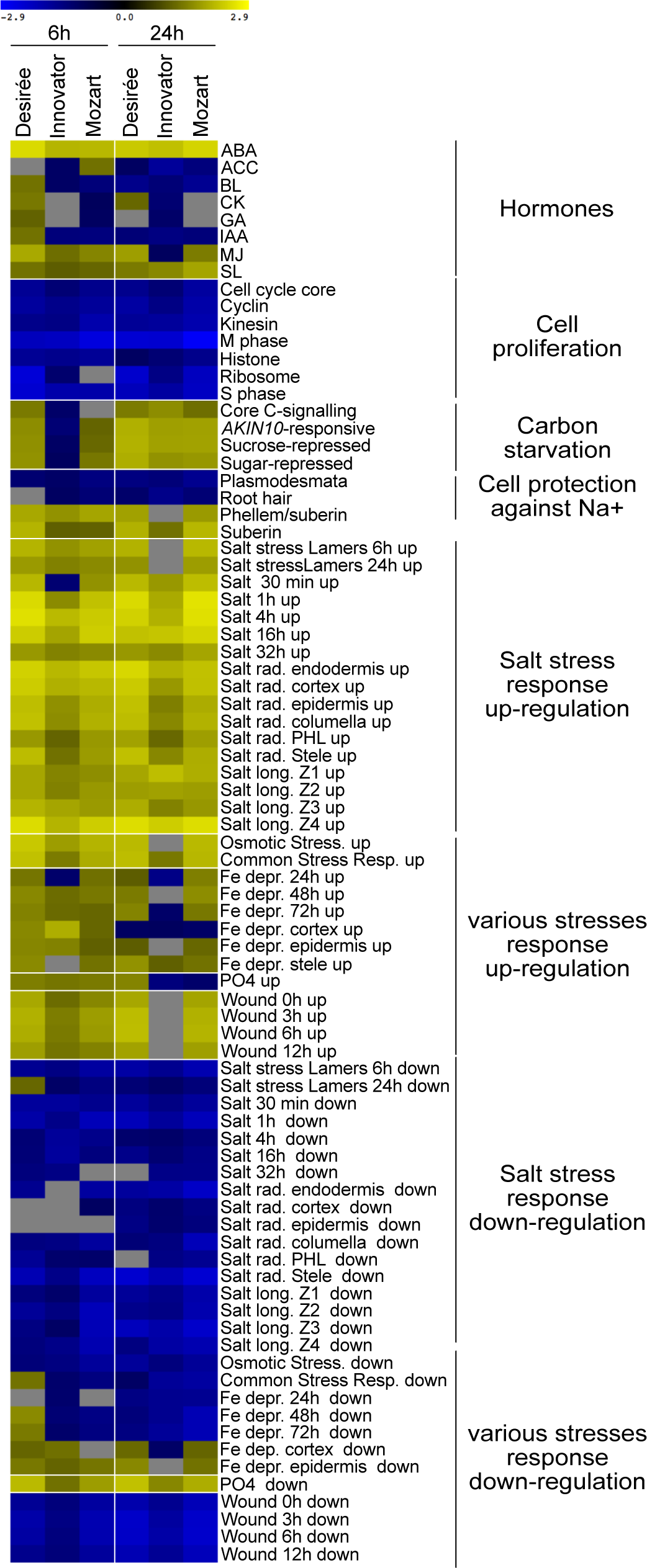
GSEA results of the salt stress response in the three cultivars. Gene Set Enrichment Analysis (GSEA) results displaying the normalized enrichment score (NES). Positive NES values (yellow) indicate gene sets which are overrepresented among induced genes and Negative NES values (blue) indicate those which overrepresented among the repressed genes. Null values indicate gene sets not significantly overrepresented.

**Fig. S3.**
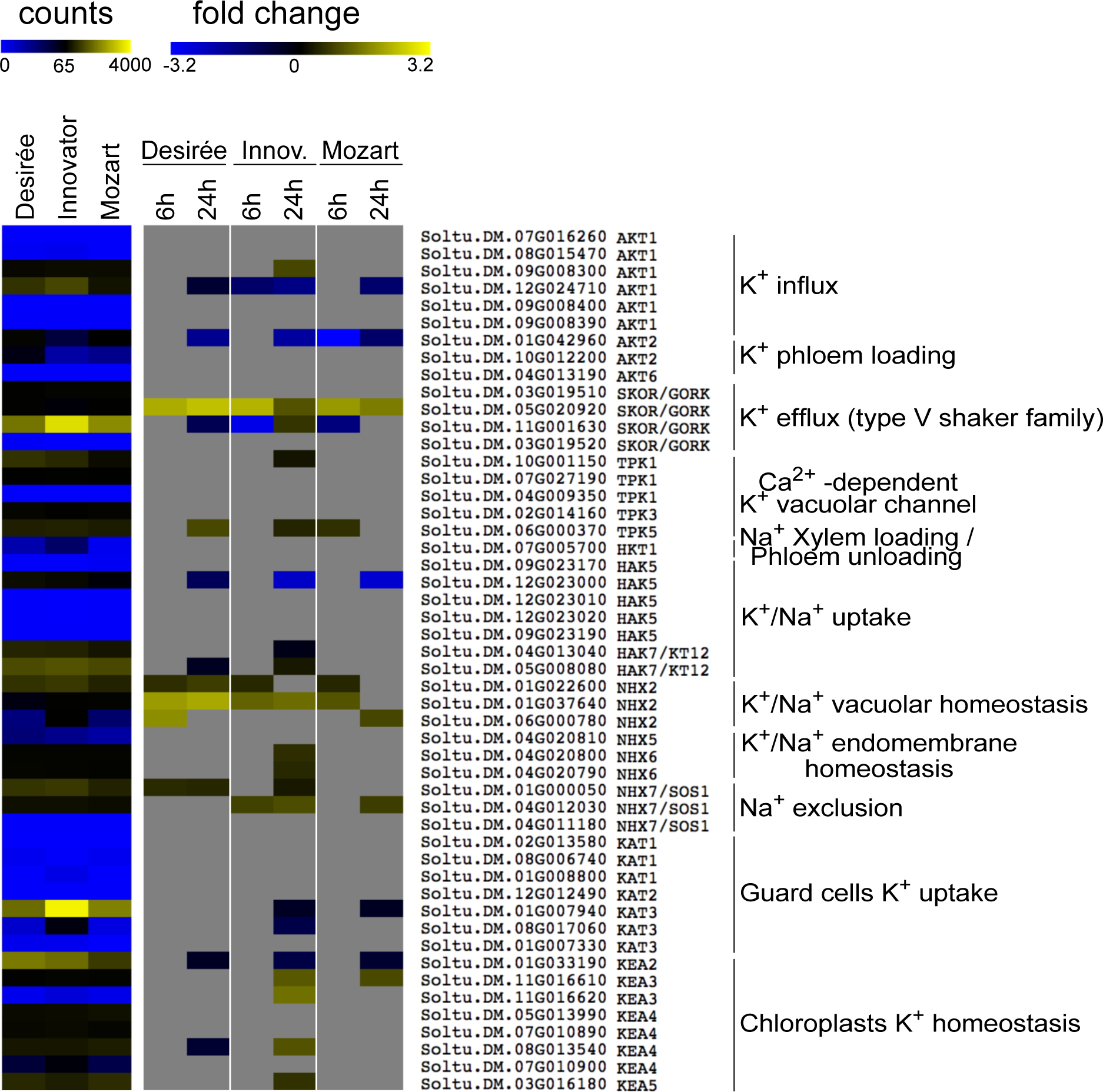
Heatmaps of the Na+/K+ channels/transporter expression pattern in roots. Heatmap on the left shows the normalized counts values at 6h in control conditions. Heatmap on the right shows the log2 fold changed values of Control versus NaCl conditions of each cultivar at a specific time point.

**Fig. S4.**
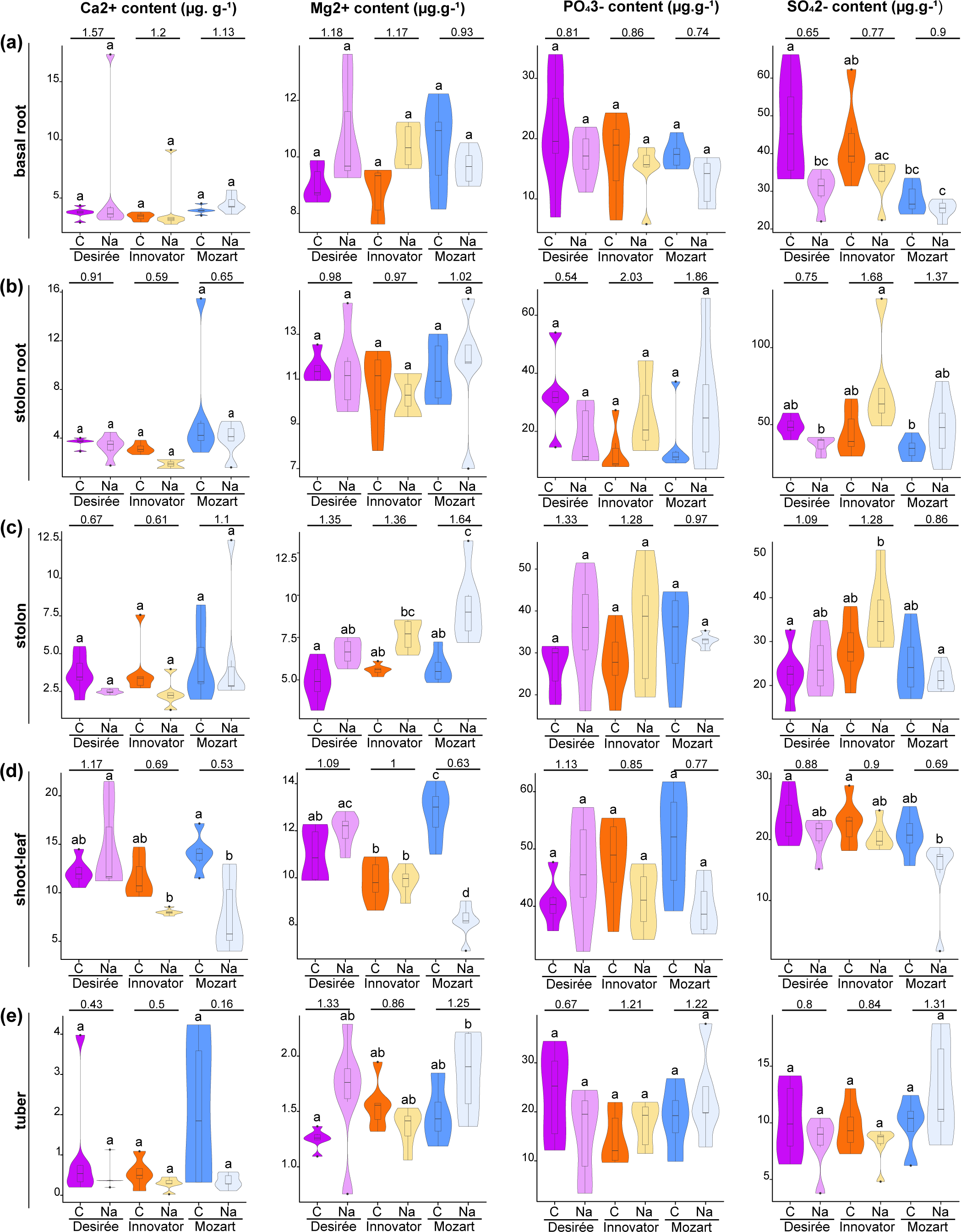
Ca2+, Mg2+, PO43- and SO42- content in salt treated Desirée, Innovator and Mozart plants. Ca2+, Mg2+, PO43- and SO42- contents in basal and stolon-node roots (a and b respectively), stolon (c), shoot land leaves tissues (d) and tubers (e). Sample size is constituted by 6 replicates, each of them from by a pool of two plants. The letters designate significant differences among means (one-way ANOVA plus Tukey’s HSD).

**Fig. S5.**
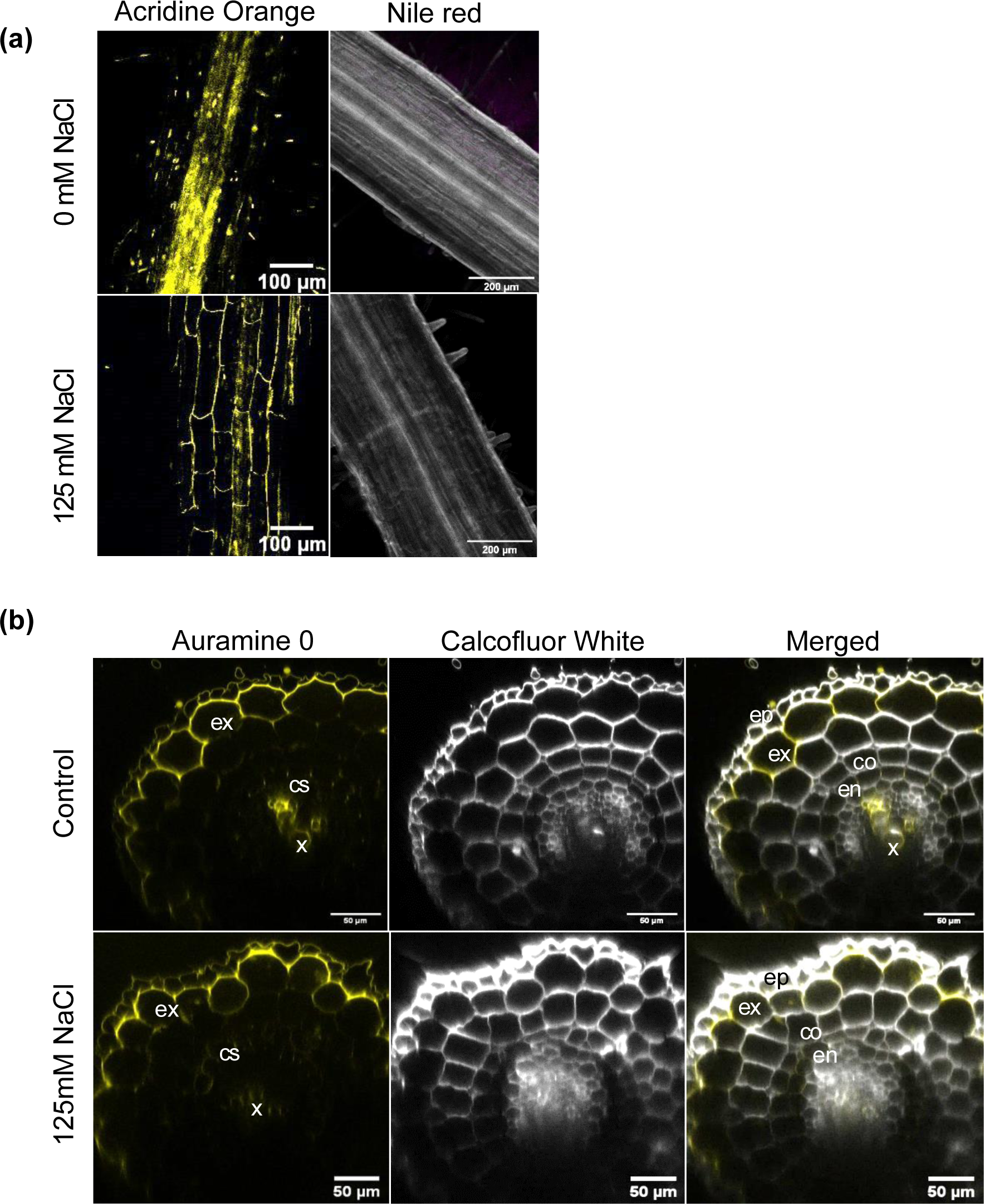
Suberin and lignin deposition in basal roots. (a) Longitudinal view of suberin and lignin deposition in potato basal roots of control and salt stressed plants (125mM). Plants were grown in in vitro conditions. Signal was observed after staining with acridine orange suggesting presence of suberin or lignin. No signal was observed after staining with Nile Red. Roots cells wall were staining by Calcofluor White (white signal). (b) Transversal view of basal root exhibiting suberin and lignin deposition. Suberin and lignin were stained by auramine O and cell wall by calcofluor white. ep indicates epidermis; ex, exodermis; co, cortex; en, endodermis; cs, Casparian Strip and x, xylem vessels.

**Method S1** RNA extraction and raw data analysis

RNA was extracted with the FavorPrep Plant Total RNA Mini Kit (FAVORGEN). DNA was digested in the column with RNase-free DNase I (Roche). The samples were sent for sequencing at Novogene (Cambridge, UK). Read quality was assessed with FASTQC (Andrews, 2010) and MULTIQC packages (Ewels et al., 2016). Trimming of low-quality reads was processed with TRIM GALORE! (Krueger et al., 2023). Reads were mapped to the transcriptome of S. tuberosum v6.1 (Pham et al., 2020) using SALMON (Patro et al., 2017). Differential Expressed Genes were obtained from pairwise comparisons using the R package DESEQ2 (Love et al., 2014).

**Method S2** ABA measurement

Ground samples were extracted with 1 ml of 10% methanol containing 100 nM stable isotope-labeled internal standards ([2H6]ABA). Abscisic acid was extracted and measured using a modified protocol (Floková et al., 2014) where a StrataX spe- column 30mg/3ml (Phenomenex) was used. Solvents were removed using a speed vacuum system (ThermoSavant). Liquid chromatography-tandem mass spectroscopy was used for the detection and quantification of ABA. Sample residues were dissolved in 100 µL acetonitrile /water (20:80 v/v) and filtered using a 0.2 µm nylon centrifuge spin filter (BGB Analytik). A Waters XevoTQS mass spectrometer equipped with an electrospray ionization source coupled to an Acquity UPLC system (Waters) was used to quantify hormones by comparing retention times and mass transitions with standards as previously described (Schiessl et al., 2019; Gühl et al., 2021). Chromatographic separations were performed using acetonitrile/water (+ 0.1 % formic acid) on a Acquity UPLC BEH C18 column (100 mm, 2.1 mm, 1.7mm Waters) at 40 °C with a flowrate of 0.25 mL/min. The column was equilibrated for 30 minutes with the solvent (acetonitrile /water (20:80 v/v) + 0.1% formic acid). For analysis, 5 µL of sample was injected, followed by an elution program in which the acetonitrile fraction linearly increased from 20% (v/v) to 70% (v/v) in 17 minutes. Between samples, the acetonitrile fraction was increased to 100% and maintained there for one minute to wash the column. Before injecting the next sample, the acetonitrile fraction was set to 20 % in one minute. A capillary voltage of 2.5 kV was used in combination with a source temperature of 150°C and a dissolution temperature of 500°C. A IntelliSmart MS Console (Waters) was used to optimize the cone voltage and multiple reaction monitoring was used for quantification (Schiessl et al., 2019). The IntelliSmart MS Console was used to set Parent-Daughter transitions. Peaks were analyzed using Targetlynx software and samples were normalized for the internal standard recovery and expressed relative to the sample fresh weight. A standard curve was used to convert peak area to pmol per mg of fresh weight.

**Method S3** Root staining for suberin and lignin quantification

3cm-long root endings were fixed in 4% PFA dissolved in phosphate buffer (PBS) for 1.5 hours in a vacuum. Roots were washed twice in PBS (10 min with gentle agitation). Afterwards roots were cleared in Clearsee (Kurihara et al., 2015) for four days in the dark. Auranine O staining was carried out by placing the root in ClearSee solution containing 0.5% Auranine O and 0.1% Calcofluor for 12 hours with gentle agitation. Nile Red staining was carried out by placing the root in ClearSee solution containing 0.05 % Nile Red and 0.1% calcofluor. Acridine Orange staining was performed by placing the root in 1µM Acridine Orange solution for 12hours as described in Li & Reeve (2004). Roots were washed twice in ClearSee solution for 30 minutes. Fluorol Yellow 088 was performed by fixing the root in 20% methanol and 4% hydrochloric acid for 20 min in 60°C. Roots were washed afterwards in 7%NaOH and 60% ethanol for 15 min at room temperature. Rehydration was done by successive EthOH baths of 5 minutes, decreasing from 40% EthOH to 20% EthOH and lastly 10% EthOH. Up next, roots were stained in a 0.01% Yellow 88 (in methanol) and were shaken for 3 days at (350 rpm) in the dark. Roots received a counterstaining with a 0.05% Aniline Blue solution (0.5% in Milli-Q water), for about an hour and washed three times in Milli-Q water.

The samples were mounted on slides containing a drop of Clearsee solution. Subsequently, the slides were put under a Leica TCS SP5 HyD confocal microscope using a HC PL FLUOTAR 10x/0.30 DRY objective to examine areas 2 cm away from the root tip. The following settings were applied: XY=512x512; 400HZ; 1,00 AU pinhole and smart gain + smart offset: disabled. A HyD1 detector was being used on counting mode. For Calcofluor White imaging, an OPSL 405 laser (strength 2.0%) was used for the excitation, with a 418-458 nm detection range. Auramine O, Acridine Orange and Yellow088 signals were visualised with an OPSL 488 laser (strength 15%) and detected with a range band from 495-554 nm. The detection range for Nile red was set from 580-620nm.

**Method S4** NaCl-induced root K^+^ flux analysis using MIFE

Seed tubers of Desirée, Mozart and Innovator were sprouted in vermiculite in a greenhouse at 22/18°C day/night set point temperatures, 40% relative humidity, a 14/10h light/dark cycle at a minimum PPFD of 170 µmol m^-2^ s^-1^ by adding a 50% Hoagland nutrient solution with a 1:1 NO3/NH4 ratio as N-source. At a stem size of about 10 cm, sprouts were carefully removed from the tuber and transferred to 30 L containers with an aerated 50% Hoagland nutrient solution. After 7 days of acclimation to the hydroponic culture, the NaCl concentration in half of the containers was gradually increased to 75 mM in three days, whereas sprouts in the other half of the containers were kept under control conditions without NaCl. Subsequently after 2 weeks, in which the nutrient solution was refreshed after the first week, primary laterals of the basal roots with a total length of 2-3 cm were collected for K^+^ flux analyses using the microelectrode ion flux estimation (MIFE) system (University of Tasmania, Hobart, Australia) similar as described by Staal et al. (2011). The microelectrodes were made by pulling borosilicate glass capillaries (0.86 mm internal diameter; Harvard Apparatus, Cambridge, UK) in a vertical pipette puller (L/M-3P-A, List Medical Electronics, Darmstadt, Germany) and transferred to a stove for 4 h or overnight at 250 °C. They were salinized with 60 μL chlorotributylsilane 7 (Sigma-Aldrich Chemie GmbH, St. Louis, WA, USA) and kept in the stove for 45 min to make them hydrophobic. K^+^-selective electrodes were backfilled with 200 mM KCl and front filled with potassium Ionophore I – Cocktail A (Sigma-Aldrich), then inserted in a holder filled with 200 mM KCl and calibrated with 0.25, 0.5 and 1,0 mM KCl. Only electrodes with a Nernst slope between -50 and -59 mV and a correlation coefficient of at least 0.999 were used. Prior to the measurements, the roots were mounted in a petri dish using bee wax and a piece of glass capillary and incubated for 1h in a bath solution containing 0.5 mM CaCl2 and 0.5 mM KCl. To determine the optimal distance from the root tip for measuring K^+^ fluxes, flux profiles along the distal 2 mm of the root of all three cultivars were taken with 250 µm increments from the root tip. K^+^ fluxes were measured for approximately two minutes at each position in a control bath solution without NaCl and after 1 h of incubation in a 75 mM NaCl-containing bath solution. The optimal distances for stable K^+^ flux analyses were established at 250, 500 and 500 µm from the root tip for Mozart, Desirée and Innovator, respectively, which are all located at the flanks of the net K^+^ efflux peak. At these distances, K^+^ fluxes from lateral roots of potato sprouts -both control and NaCl acclimated- were recorded in the bath solution without NaCl to record the baseline K^+^ flux values. After 5 min, 75 mM NaCl was added, and the NaCl-induced K^+^ flux was recorded for a subsequent 25 min. Five biological replicates per treatment and genotype were used. Data represent the mean, error bars the standard deviation.

